# Vortex chip incorporating an orthogonal turn for size-based isolation of circulating cells

**DOI:** 10.1101/2020.08.15.251991

**Authors:** Navya Rastogi, Pranjal Seth, Ramray Bhat, Prosenjit Sen

## Abstract

Label-free separation of rare cells (e.g. circulating tumor cells (CTCs)) based on their size is attractive due to its wider applicability, simpler sample preparation, faster turnaround, better efficiency and higher purity. Amongst cognate protocols for the same, vortex-trapping based techniques offer high throughput but operate at high flow velocities where the resulting hydrodynamic shear stress is likely to damage cells and compromise their viability for subsequent assays. We present here an orthogonal vortex chip which can carry out size-differentiated trapping at significantly lower (38% of previously reported) flow velocities. Fluid flowing through the chip is constrained to exit the trapping chamber at right angles to that of its entry. Such a flow configuration leads to the formation of vortex in the chamber. Above a critical flow velocity, larger particles are trapped in the vortex whereas smaller particles get ejected with the flow: we call this phenomenon the turn-effect. We have characterized the critical velocities for trapping of cells and particles of different sizes on chips with distinct entry-exit configurations. Optimal architectures for stable vortex trapping at low flow velocities are identified. We explain how shear-gradient lift, centrifugal and Dean flow drag forces contribute to the turn-effect by acting on cells which pushes them into specific vortices in a size- and velocity-dependent fashion. Finally, we demonstrate selective trapping of human breast cancer cells mixed with whole blood at low-concentration. Our findings suggest that the device shows promise for the gentle isolation of rare cells from blood.

## Introduction

The isolation of a specific cellular niche from heterogeneous populations has been an enduring topic of intense investigation at the interface of biology and microfluidics^1-3^. This is especially relevant in the context of the separation of circulating tumor cells (CTCs) from the blood of patients afflicted with cancer. CTC detection can be used as a diagnostic assay^4^, a method to determine the efficacy of treatment^5^ and to indicate the relapse of cancer^6, 7^. The ability to culture the captured patient-derived CTCs^8^ would allow us to further investigate disease progression and therapeutic outcomes in greater detail and on a personalized level^9^.

Existing technologies carry out separation by either inducing external forces (active separation) or by using intrinsic hydrodynamic forces (passive separation). Active separation methods include dielectrophoresis^10^, magnetophoresis^11^, acoustophoresis^12^, etc., where the flow direction of the selected particles is controlled or modified using an external force. Drag-induced primary motion of particles is diverted by the external forces being invoked. With these external forces typically in the order of pN, the sample processing rate for such techniques is low, with volume flow rates of about 100 µL/min and particle translation speed in the range of 100 µm/s. This ultimately results in a limited throughput. Moreover, these techniques often involve biomarkers^13-18^ and labeling agents^19^ which may be incompatible with strategies for further culture of these cells and clinical applications. Additionally, active separation strategies often fail to isolate the entire population of target cells^20^ due to the inherent heterogeneity in the expression of markers and membrane proteins which are often the ligands for labeling agents and enriching methodologies^21, 22^.

Passive separation on the other hand is carried out by simply controlling the hydrodynamic properties of flow. These include pinched flow fractionation^23^, Dean flow fractionation^24^, deterministic lateral displacement^25^, microfiltration^26^, inertial microfluidics^27-29^, etc. These techniques separate and concentrate cells based on the difference in their inherent physical properties such as size, deformability, etc. Inertial microfluidics in particular provides a high-throughput, whereby large sample volumes can be processed within a short time interval. Operating at flow rates of up to 5 mL/min, this advantage is especially useful for processing high volumes of bodily fluids such as blood and ascites for tumor cells. Owing to the rarity of CTCs, the capability to quickly process comparatively large volumes is a practical necessity. Shorter processing times are also known to result in lower death rates of CTCs which enhances their usability for future assays^30, 31^.

In the flow regime where the inertial effects become pronounced, a separating Stokes flow is seen for flows around bodies placed in a uniform flow or flows in a channel with sudden expansion^32^. These are known as Moffat eddies^33^, which can grow into fully developed microscale laminar vortices as the Reynolds number of the flow increases. Such vortices combined with inertial focusing at sufficiently high Reynolds number have been strategically utilized by Di Carlo and coworkers to carry out size-selective isolation of cells^20, 29, 34, 35^. However, these vortex-trapping techniques are able to operate only at high flow-velocities of up to 4 m/s, where the resulting hydrodynamic shear stress is likely to damage the cells and compromise their viability^36-38^ for downstream applications.

In this paper, we describe the design and characteristics of a passive and continuous microfluidic device that is capable of size-based trapping and separation of rare cells from a heterogeneous suspension. The aim of the new design is to achieve trapping at lower flow velocities. Fig. 1A shows a generalized schematic for this vortex chip with orthogonal arrangements of entry and exit for fluid flow. Within the trapping chamber, configuration-specific formation and growth of microscale-vortex with flow velocity is observed. Above a critical velocity, the vortex selectively entraps larger beads and cells, since they are ejected out of the flow owing to the ‘turn effect’. This turn effect is due to a combination of physical forces arising in the fluid flow trajectory due to the sharp turn resulting from the orthogonal entry-exit configuration. Meanwhile, smaller red blood cells predominantly escape the trapping chamber with the flow, thus enabling separation of the rare larger cells. This device operates at flow-velocities that are up to 38% of the previously reported vortex-trapping methods^20, 29, 34, 35, 39, 40^, which should improve the viability of cells for culture-based applications. Further, lower velocities imply lower operational pressures, which simplifies device fabrication and operation. Exploiting some of the unique features of inertial^41, 42^ and non-linear^43^ microfluidics, we present here a simple and practical size-differentiated separation technique which could have broad applicability.

**Figure 1:**
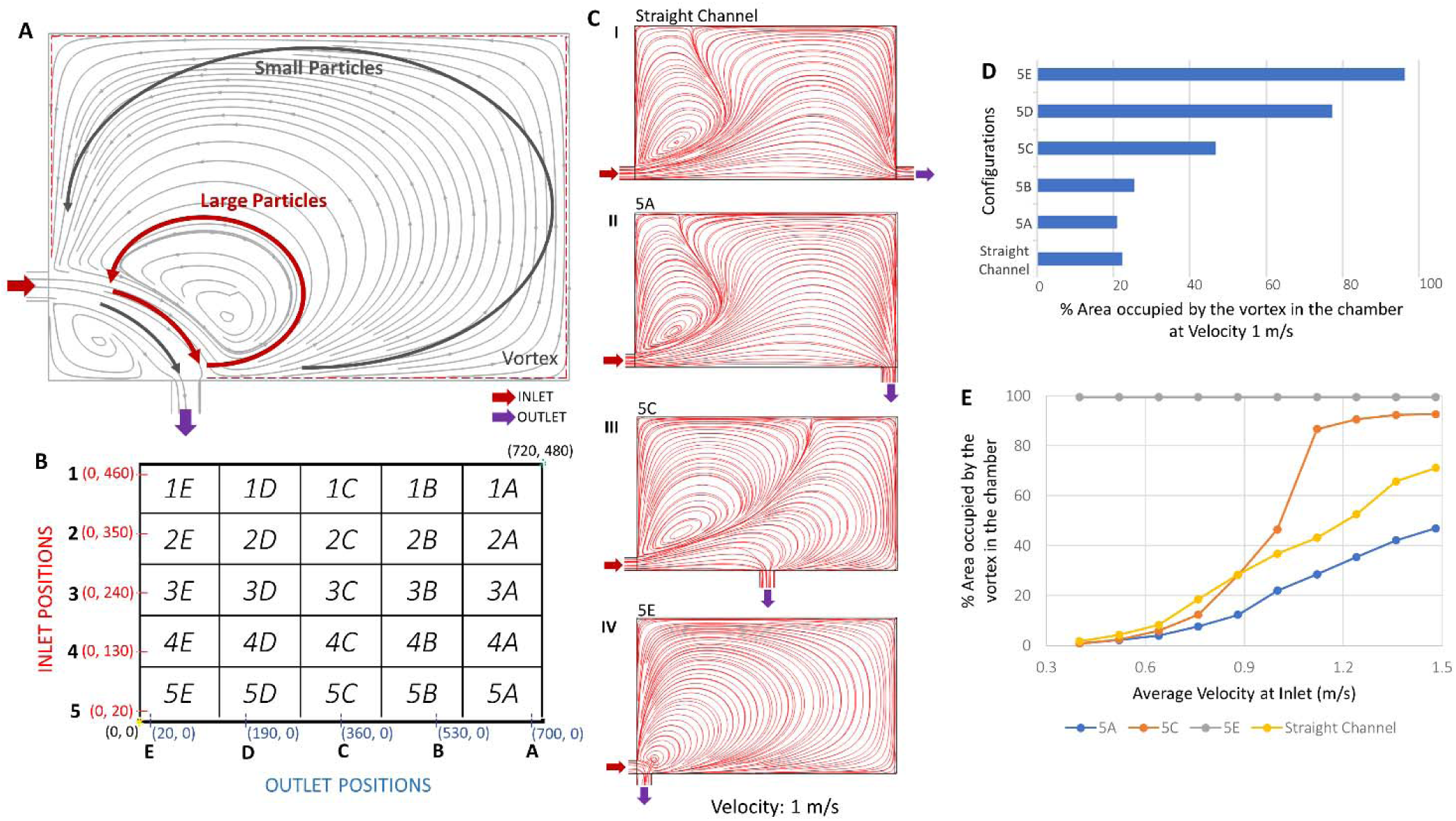
**A**. A generalized schematic of the orthogonal vortex chip which carries out size-differentiated vortex trapping. Large particles get trapped in the inner orbits close to the vortex core, while smaller particles just flow along the stream or circulate in the unsteady outer region of the vortex and pass by without getting trapped. **B.** A coordinate-map of the orthogonal vortex chip variations along with the naming nomenclature. The x-y plane is shown, and the origin is taken to be at the bottom-left corner of the reservoir. There are five different positions for both inlet and outlet, resulting in 5×5=25 total combinations. An inlet at label ‘3’ would have its midpoint at (0, 240), and an outlet at label ‘C’ would have its midpoint at (360, 0). Such a design would be called ‘3C’ device. It should be noted that a reservoir has only one inlet and one outlet at a time. All coordinates are in microns. **C.** (I-IV) Geometry with inlet and outlet closer in proximity to each other forms larger vortices for the same flow velocity. Shown here are COMSOL simulation flow-streamlines for configurations: straight channel (no turn) (I), 5A (II), 5C (III), and 5E (IV) for average velocity U = 1m/s and Re = 46. **D.** Percentage of area occupied by the vortex in configurations straight channel and 5A-E. **E.** Change in the size of the vortex with increasing flow velocity for configurations: straight channel, 5A, 5C and 5E. Configurations with an outlet located closer to the inlet exhibit saturation with a fully developed vortex that occupies most of the reservoir area at lower flow velocities.

## Results and discussion

### Simulation Results

Previous efforts at vortex based size-specific cell trapping utilized straight flows at high Reynolds numbers ^20, 29, 34, 35, 39, 40^.

We began by asking whether introducing an orthogonal bend in the flow trajectory might allow formation of a large vortex suitable for stably trapping particles and cells at lower flow rates. We used COMSOL Multiphysics to simulate fluid flows through such designs. 3D flow simulations were carried out and Navier-Stokes equations for incompressible flow were utilized in the Laminar Flow module. No-slip boundary condition was used at the walls and zero pressure boundary condition was used at the outlet. Flow conditions were varied by changing the average flow velocity at the inlet (See Fig. S1). The trapping chamber dimension was kept the same (480 µm × 720 µm) as in previous efforts by other groups^20, 29, 34, 35^ in order to enable comparison with those results.

Several distinct variations of this orthogonal configuration based on the location of inlet and outlet were studied. Fig. 1B provides a detailed description of the nomenclature system used for identifying device geometries. Flow simulations were performed for these configurations with the objective of generating maximally sized vortices at the lowest flow velocities. Fig. 1C shows flow streamlines from COMSOL simulations for selected configurations suggesting that as the inlet and outlet are brought closer in proximity to each other, the area covered by the vortex increases for the same average flow velocity of 1 m/s. The percentage area covered by the vortex for these configurations was quantified using ImageJ and is shown in Fig. 1D. Configurations with outlet placed closer to inlet are observed to yield a larger vortex. Fig. 1E shows the change in area occupied by the vortex in these geometries for increasing flow velocity. Configurations with inlet and outlet placed closer to each other are observed to attain a saturated state, where the vortex occupies most of the reservoir area, at lower flow velocities. The simulations therefore indicate that certain orthogonal flow designs produce larger vortices at lower velocities.

### Flow and Trapping Characterization

Next, we sought to experimentally verify our modeling results. We further wanted to characterize the minimum velocity required for trapping particles of a given size. Microfluidic devices were fabricated with PDMS using soft lithography as described in Methods. A suspension of microparticles (polystyrene beads of 12 µm and 15 µm diameter) was flowed in order to visualize the flow streams, verify the presence of the vortices, study their shape, characterize the velocities required for trapping and examine the particle behavior.

We observed that the trajectory of particles in these microparticle-laden flows depends on both particle size and flow velocities, as shown in Fig. 2A for design 4C. At lower velocities, the particles follow the streamlines and exit the chamber with the flow. For instance, 20 µm and 15 µm beads do not get trapped up to 1.12 m/s and 1.24 m/s respectively. Above a critical velocity, the particles start escaping the flow and get trapped in the vortex, which then appears as an enclosed recirculating trajectory. We define this minimum flow velocity at which particles of a given size get trapped in the vortex as the *trapping-velocity;* and the recirculating path itself is defined as the *trapping-orbital.* Experimentally observed trapping-velocity for 20 µm and 15 µm beads in design 4C is 1.24 m/s and 1.36 m/s respectively. Increasing flow velocity further also reveals multiple stable orbits for 15 µm beads at 1.36 m/s. Thus, for a fixed flow velocity, different-sized particles exhibit their own unique trajectories in the reservoir with larger particles getting trapped at lower velocities. Moreover, while COMSOL simulations confirm the formation of vortex even at low velocities, vortex trapping is observed only above certain flow velocities which is specific for a given particle size. This implies that vortex formation in itself doesn’t necessarily lead to trapping of particles and a nuanced effect of inertial forces is at work here.

**Figure 2:**
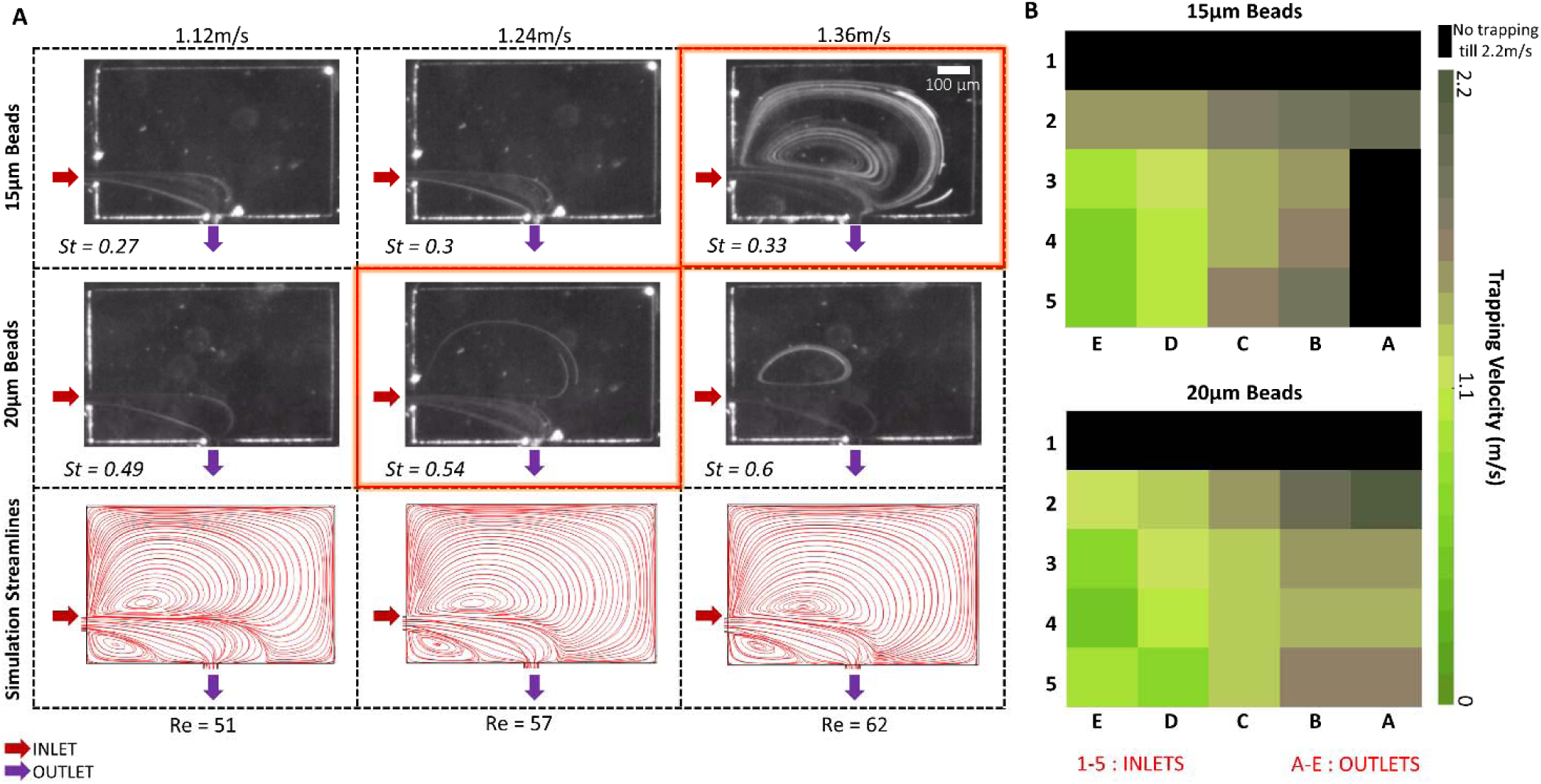
**A**. (Top two rows) Flows laden with microparticles behave differently for various flow velocities and particle sizes, where different-sized particles exhibit their own unique trajectories in the reservoir. A particle is said to be trapped in the vortex if an enclosed recirculating trajectory is observed (highlighted box). (Bottom row) COMSOL simulation streamlines for the same conditions, which shows that the presence of vortex doesn’t necessarily imply the entrapment of particles. Mentioned here is respective average flow velocity, particle size, Reynolds number and Stokes number for the different cases. Scale bar represents 100 µm. Arrows indicate inlet and outlet flow direction**. B.** Trapping velocity of particles (15µm and 20µm) for all the 25 different inlet-outlet designs.

For each inlet-outlet configuration, a different trapping velocity specific to each particle size was observed. Trapping-velocity for 15 µm and 20 µm beads was experimentally determined for all inlet-outlet configurations as shown in the 2-dimensional color-coded map in Fig. 2B. Velocity- and size-specific behavior pertaining to trapping-velocity described above for design 4C was observed to be consistent for other inlet and outlet configurations as well. In general, as the inlet and outlet are brought close to each other, we observe that the trapping velocity decreases. This is expected as the turn-radius decreases when the inlet and outlet come closer. *Turn-radius* is defined as the effective radius of the circular path taken by flow at the turn (see Fig. S2A). As explained in the later section describing the turn effect, a smaller turn-radius enhances the inertial effects responsible for the ejection of particles from the flow stream.

However, the distribution of trapping velocity in Fig. 2B is neither monotonic nor symmetric. For example, we observe that for the closest inlet-outlet configuration (design 5E), the trapping velocity for the 20 µm beads is higher than the neighboring configurations. This is possibly due to the extremely short path of the bead in the trapping chamber. Due to the shorter flow path within the chamber, larger inertial forces are required to obtain the requisite deflection for the particle to leave the flow stream. Further, at the velocities used in these studies, the flow does not follow the shortest path between the inlet and outlet for some of the designs, rather it takes an arched trajectory (as seen in Fig. S2A). Hence, the turn-radius has a non-trivial relationship with respect to inlet-outlet position. We believe that these are the underlying reasons for the observed deviations in the distribution of trapping velocity.

Finally, for the same flow velocity and bead size, a variation in the inlet-outlet configuration also led to a change in the size and position of the trapping-orbital around the vortex (Fig. S3). Considering steric effects, a purely geometric argument implies that the number of beads that could be trapped within a vortex also changes with the inlet-outlet configuration. Even though trapping occurs at lower velocities for the closer inlet-outlet configurations, the obtained trapping-orbitals were small. Steric effects should limit the number of particles that can occupy these orbitals. Larger trapping-orbitals were observed when the distance between the inlet and outlet was increased. These larger orbitals should be able to accommodate a greater number of particles before the steric effects start to play a role. However, at the same time, placing the inlet and outlet far apart required higher trapping velocity and resulted in unsteady trapping. Unsteady trapping is characterized by two observations. Firstly, the particles are observed to leave the main flow stream and enter the chamber. However, they are unable to get into a steady orbit and exit the chamber without getting trapped. Secondly, in cases where the particles are trapped, one or more particles in the trapping-orbital were observed to leave the trap and exit the chamber after some time, possibly due to small flow rate fluctuations (see Fig. S2A). This suggests an optimum zone whereby stable trapping could be accomplished at low velocities. Hence, out of the 25 possible configurations, designs 3D, 3E, 4D and 4E were chosen (Fig. S2B) for further studies as these exhibited stable particle trapping at the lowest possible velocities. The suitability of these designs is addressed again in the section describing the turn effect.

We also conducted high-speed videographic studies to observe the trapped particles in these four designs that were deemed optimal. A suspension of particles with reduced concentration was used in order to observe individual particle behavior in the reservoir. A steady entrapment was seen in most cases, but there were instances where the particles already trapped in the vortex would get ‘knocked out’ (See Fig. 3A and Movie M1) due to a new incoming particle. Few particles were also seen to enter and exit the vortex area without getting trapped in an occupied orbital. Both of the above observations suggest that steric effects play a role in limiting the number of particles that every trapping-orbital can accommodate. The trapping-orbitals serve as “attractor-zones”, where they exhibit steady entrapment for the particles. Separate stable orbitals would exist corresponding to every particle size at a given flow rate. Also, multiple attractor-zones or trapping-orbitals may coexist within the vortex as seen in Fig. 2 for 15 µm beads at flow velocity 1.36 m/s. Particles were also observed to be trapped in recirculating trajectories in different vertical planes (Fig 3A(i)). This might be due to the existence of distinct flow profiles in planes at different depths (Fig. S4) in the reservoir because of which multiple stable orbitals are possible.

**Figure 3:**
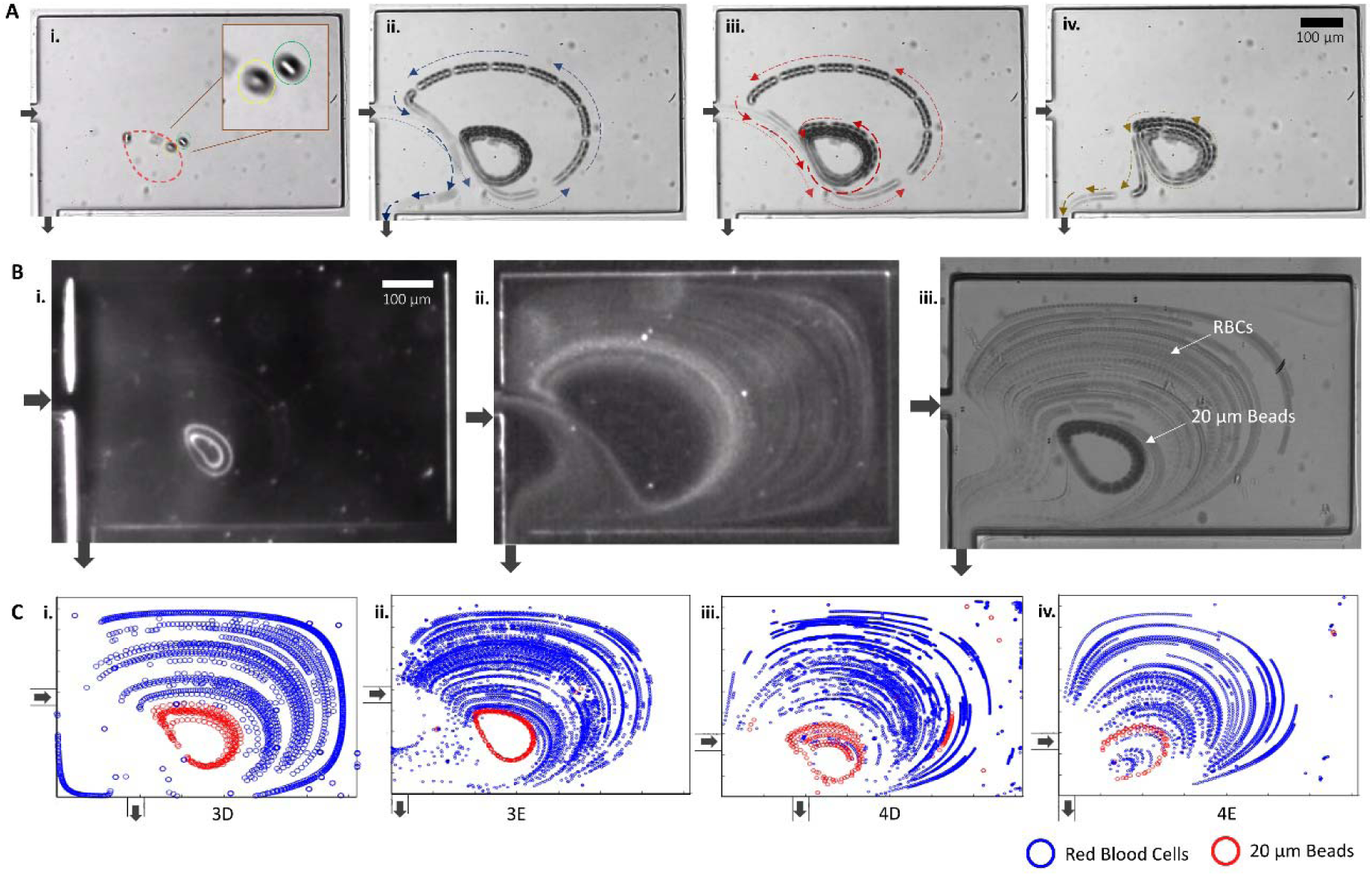
**A** A still and an overlay of frames when 20µm beads are flowed in the device. Images from the high-speed studies showing **i**. Beads are observed to be trapped in different focal planes, indicating that the captured beads are recirculating in planes at different depths **ii.** A bead enters the reservoir and exits after traversing through the outer orbital without getting trapped. This confirms that the outer orbitals of the vortex do not enable stable trapping. **iii.** A new bead gets trapped with the already trapped beads when it enters the reservoir **iv.** A trapped bead gets knocked out of the attractor zone. **B** When different-sized particles are flown individually: **i**. 20 µm beads get trapped in the steady trapping orbital close to the core of the vortex. St_Large-Beads_ ≈ 0.5. **ii**. RBCs pass by undisturbed or follow the unsteady orbits in the outermost zone of the vortex. St_RBCs_ ≈ 0.05. **iii.** An overlay of frames when different sized particles are flown together as a heterogenous mixture. 20 µm beads get steadily trapped in the vortices, as indicated by the enclosed recirculating trajectory, while RBCs just pass by. **C i.** Corresponding size-labeled scatterplot of the tracked particles and cells inside the microvortex flow in the reservoir, for the *3D* design. Similar scatter plots are obtained for **ii**. *3E* **iii**. *4D* **iv**. *4E*. Scale bar represents 100 µm. All flows are at U = 1.3m/s.

### Trapping Induced Separation of Cells/Particles

We next asked whether beads of large sizes would continue to get trapped within vortices upon flowing in a heterogeneous suspension that also consisted of red blood cells (RBCs) as smaller particles. As discussed before, particle-particle or particle-cell interaction cannot be neglected. Hence, there is always a possibility that larger but rare particles in the trapping-orbital are ‘knocked-out’ by one of the numerous smaller cells. Successful trapping under this scenario necessitates that the trapping-orbital of the larger particle is distinct from the path taken by the smaller cells (RBCs).

To understand the trapping behavior, beads and diluted whole blood were flowed separately at different velocities. Above a critical velocity, the larger beads (20 µm) were trapped in the stable orbitals close to the vortex core as seen in Fig. 3B(i). On the other hand, blood cells when flowed at the same velocity through the same design were observed to exit it without entering the vortex, or merely exited after traversing the outermost orbitals of the vortex as seen in Fig. 3B(ii). The exclusivity of the trapping-orbital and the path taken by RBCs provided a flow regime where trapping-induced separation or concentration enhancement would be possible.

When heterogeneous samples consisting of both blood cells and beads were flowed together (see Methods for details of sample preparation), they retained their individual behaviors (Fig. 3B(iii) and Movie M2). This was also confirmed through the scatterplots generated using the high-speed camera videos, capturing the flow of heterogenous mixture of 20 µm beads and RBCs (Fig. 3C). Earlier reports have confirmed that the outermost region of microfluidic vortices tend to be unstable allowing particles to easily escape the vortex, whereas particles get stably trapped in the innermost orbits^39^. This allows us to execute a vortex-trapping induced separation of larger particles or cells. We sought to explain this phenomenon of size-specific entrapment and localization within vortices in the reservoir with orthogonal inlet and outlet by investigating the forces acting on the particle and measuring its Stokes number.

### Turn-Effect

The observed turn-effect is a result of multiple inertial forces being exerted on the particle in flow. In this section, we discuss these forces and their individual contribution to the phenomenon. A schematic representation of all the forces is shown in Fig. 4A. While flowing in the channel, particles experience hydrodynamic inertial focusing as they travel downstream, being subjected to a shear-gradient lift force directed towards the channel wall and an opposing wall-effect lift force directed towards the channel centerline^44^. Using scaling arguments, the resultant lift force has been described as^45^:

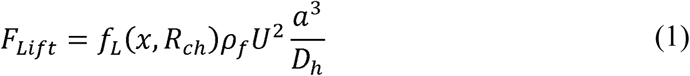

where *ρ*_*f*_ is the density of fluid, *a* is the particle diameter,*U* is the flow velocity,*f*_*L*_ is the lift coefficient which is a function of particle’s lateral position in the channel *x* and channel Reynolds number *R*_*ch*_, and *D*_*h*_ ithe hydraulic diameter of the channel defined in terms of width *W* and height *H* as *D*_*h*_ = 2*WH* /(*W*+*H*) When this focused particle enters the reservoir, the neighboring microchannel wall is no longer in its vicinity and the hydrodynamic interaction with the wall disappears^41^. In the absence of the wall-lift force, the remaining shear-gradient lift pushes the particle away from the centerline flow stream directing it down the shear-gradient towards the center of the trapping vortex^40^. (See Fig. S5)

**Figure 4:**
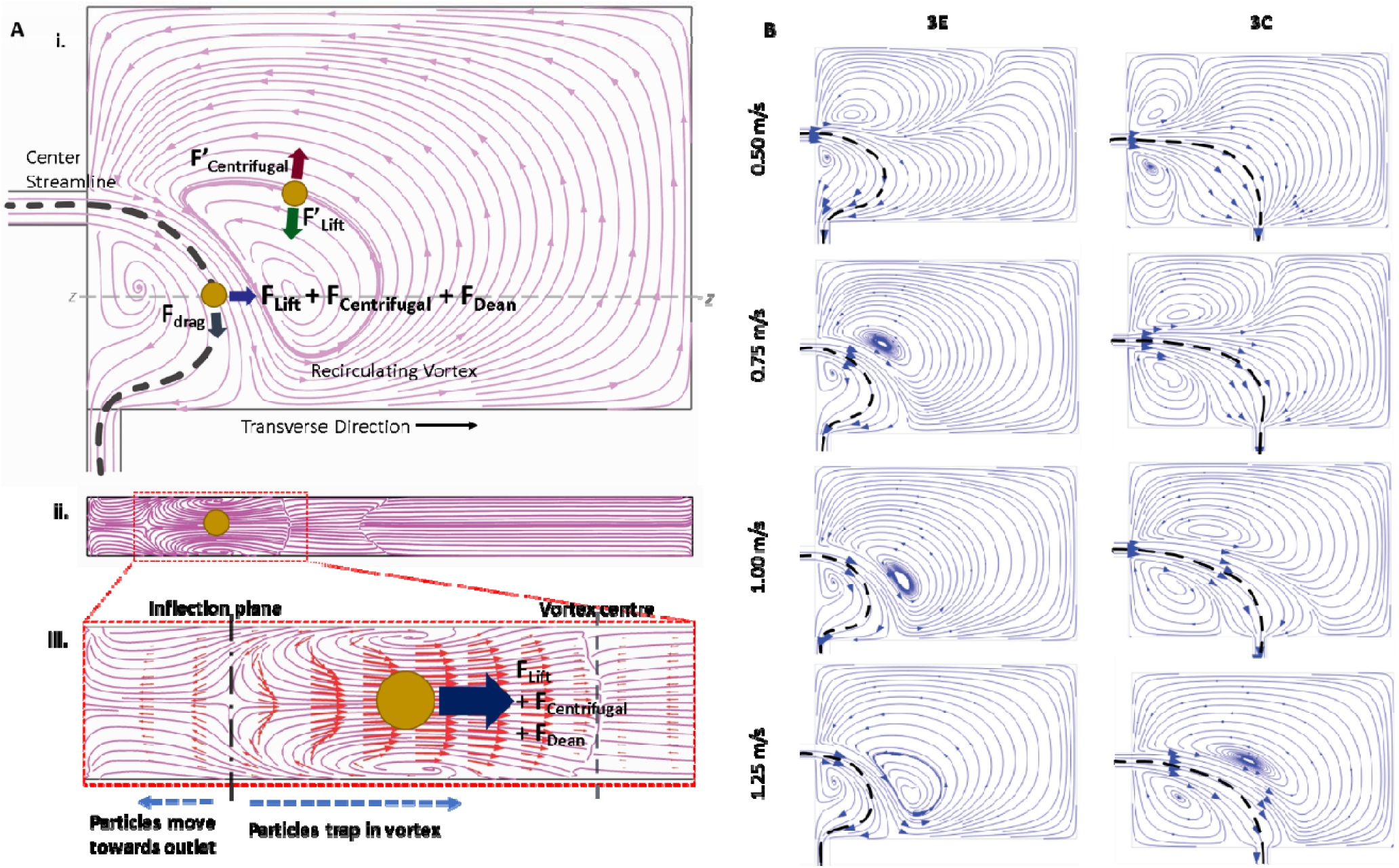
**A.** i. Sketch of the trapping reservoir in the XY plane showing the particle trajectory and the forces that act on it while passing through the turn and after it is trapped in the vortex inside the reservoir.. **ii**. A XZ plane is cut along the Z-Z line in Fig. 4A(i), **iii.** Forces on particle shown in the XZ plane, along with velocity streamlines and arrow vectors from COMSOL simulations. The streamlines confirm the presence of Dean vortices, while the velocity vectors indicate the direction of force acting on the particle. **B.** COMSOL Simulations showing flow streamlines along with approximate center streamline in designs 3E and 3C at different velocities comparing the vortex sizes and r_turn_.

Another effect arises due to the higher density of the particles in comparison to the surrounding fluid. The particle experiences centrifugal force due to the curvilinear trajectory which imparts a transverse motion away from the centerline flow stream towards the trapping vortex. This centrifugal force on the particle can be written as^46^:

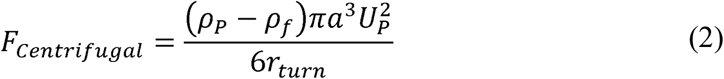

where *ρ*_*p*_ is the density of the particle, *U*_*P*_ is the particle velocity, *r*_*turn*_ is the turn-radius of the particle’s curvilinear trajectory (see Fig. S2A). This force increases with reduction of the turn-radius (*F*_*Centrifugal*_ ∝ 1/*r*_*turn*_)

The curvilinear path of the streamlines also gives rise to transverse Dean flows^47, 48^. Fig. 4A(ii) and 4A(iii) show the secondary flows in the channel cross-section. It is important to note that unlike curved channels, here the curvilinear flow path is not bounded by the channel walls. Hence a single cross-section does not capture the transverse plane for flow at different positions. We are however able to observe the plane of inflection. On the left of the plane of inflection the transverse flows drag the particle towards the outlet. However, on the right, the particles are strongly dragged towards the vortex. Secondary flow results in an additional transverse drag force on the particles, which for conventional curved channels has been stated as^49^:

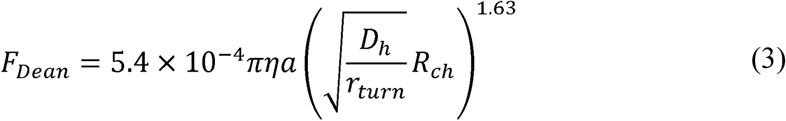

where *η* is the dynamic viscosity. The Dean drag arising here from the curvilinear flow further aids in transverse particle migration out of the centerline flow stream into the vortices. Like centrifugal force, Dean drag force also increases with decrease in the turn radius (*F*_*Dean*_ ∝ 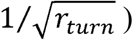. Lift force, however, is independent of the turn-radius and hence should not change with inlet-outlet configuration. Lift force (equation 1) and centrifugal force (equation 2) both are directly proportional to the cubic power of the particle diameter (∝ *a*^3^). In contrast to the other forces, Dean drag is proportional to particle diameter (∝ *a*) and hence changes more slowly with change in particle size.

The cumulative result of all the forces described above is a transverse migration force that has a complex dependency on the inlet-outlet configuration, particle diameter and velocity. In addition to these forces, the particle also experiences a drag force due to the primary flow from the inlet to the outlet:

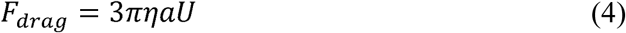

Acting in the direction of the primary flow, this drag force carries the particle along with it. Hence, the ejection of particle from the primary flow and its trapping in the vortex is the result of a competition between the primary drag and the transverse migration forces (shear-gradient lift, centrifugal and Dean drag). For a fixed particle size, at lower flow velocities, the primary drag dominates and the particles follow the flow. As the flow velocity is increased, the transverse forces increase faster than the primary drag due to *U*^2^ dependence. For a given flow condition, these forces are much more pronounced for the larger 20 µm particles compared to the smaller RBCs owing to its larger diameter. Larger particles are thus expected to migrate laterally out of the centerline flow stream and into the vortex, while smaller particles below a critical size will not migrate fast enough to cross into the vortex before passing out of the reservoir. The above estimate of forces also validates the experimental observations in which the chosen devices (designs 3D, 3E, 4D and 4E) work better with stable trapping at low velocities. For all the device configurations, the shear-gradient lift force experienced by the particle remains approximately the same. However, with the inlet and outlet closer to each other, *r*_*turn*_ is reduced (see Fig. 4B) and hence there is a larger Dean drag and centrifugal force directing the particle towards the vortex.

Furthermore, once a particle has entered the vortex, its orbit is determined by a balance^40^ of the centrifugal force, shear gradient lift force, and drag forces due to secondary flows in the transverse direction (Fig. 4A(i)). Acting in the direction of decaying shear gradient, the shear-gradient lift force is still operational in the vortex leading to a force towards the vortex center. Centrifugal force arising from the closed recirculating trajectory of the particle is directed outwards. Even though devices with inlet and outlet in immediate vicinity (designs 5D and 5E) result in smaller trapping velocities, they show unsteady trapping (see Fig. S2A). These designs have smaller trapping orbitals close to the primary flow. Smaller trapping orbital means that at equilibrium both centrifugal and lift forces are large, and also hints at steric crowding for the trapped particles. Thus, even a small change in the trapping-orbital due to fluctuations in flow velocity or disturbance due to crowding results in unstable and irregular trapping.

The previously reported linear vortex-chip depended only on the shear-gradient lift force in order to accomplish size-specific cell and particle isolation, and hence required high flow velocities in order to generate significant lift force. Notably, the incorporation of turn effect gives rise to a number of additional forces which collectively enable vortex trapping at flow velocities (1.2-1.3 m/s) that are significantly lesser compared to those of linear vortex-generating architectures.

### Particle Stokes Number

Under the influence of forces associated with the turn effect, trapping also depends on the time spent by the particle inside the reservoir and whether this time is enough for the transverse forces to sufficiently modify the particle’s trajectory. This behavior of particles in the fluid flow can be explained by considering the particle Stokes number *St*. *St* is defined as the ratio of the inertial relaxation time(*τ*_*γ*_) of the particle to the characteristic time of flow(*τ*_*f*_) ^50^:

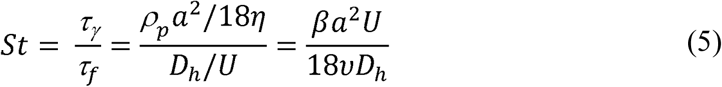

where *β* is the ratio of particle density (*ρ*_*p*_)to fluid density (*ρ*_*f*_), and υ is the kinematic viscosity. A larger *St* implies that the particle has a stronger preference to persist with its original velocity direction instead of following the accelerating fluid trajectory^50^.*St* here thus serves to be an indirect measure of the particle inertia and describes the end effect. The cells and particles eject out differentially upon introducing the abrupt orthogonal turn in the flow owing to their size dependent inertia. Larger particles (20 µm beads, *St* ≈ 0.5) bearing larger *St* shoot through the out flow-steam and transiting ahead get trapped in the vortex. Smaller particles (RBCs, *St* ≈ 0.05) meanwhile continue flowing along the stream or circulate in the unsteady outer zone and pass-by undisturbed (see Fig. S6). However, it is important to note here that while *St* offers a reasonable generic explanation of the phenomenon, it does not account for all the physical factors involved in the turn effect.

### Trapping of Cancer Cells Suspended in Whole Blood

In order to verify whether cancer cells could also be separated from a suspension in whole blood, we flowed cells from three transformed malignant breast cancer cell lines: MDA-MB-231, MCF7 and BT-549 (which show mild differences in size and shape), along with whole blood diluted in PBS. The orthogonal configuration in our device was able to effectively isolate cancer cells from other blood cells: while the former were trapped within vortices, the latter flowed out directly or after passing through outer unstable orbitals of the vortex. (Fig. 5(A-C) and See Movie *3-5*).

**Figure 5:**
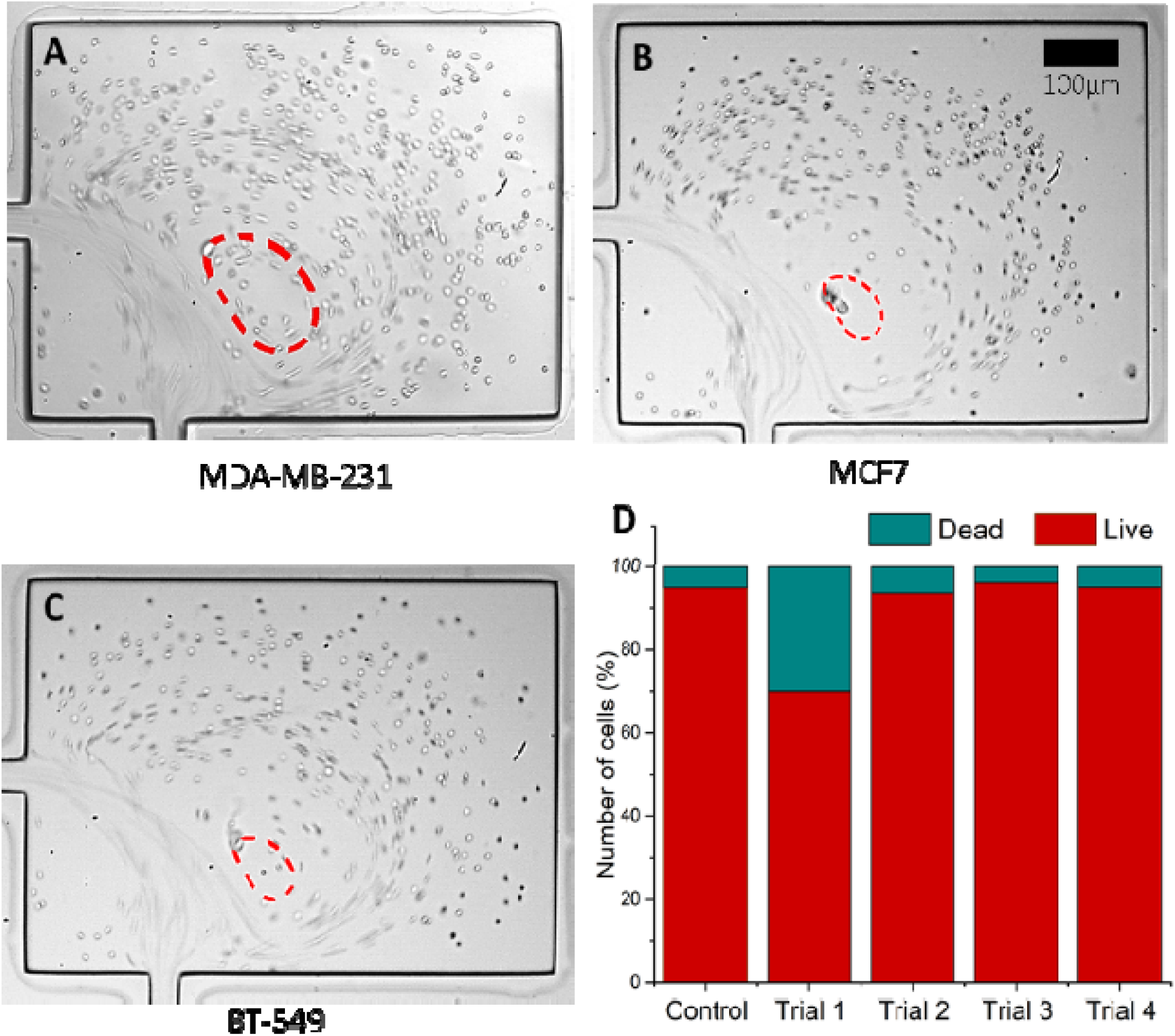
**A-C** Trajectory of 3 types of cancer cells (MDA-MB-231, MCF-7 and BT-549) suspended in whole blood when flowed through the orthogonal chip. Scale bar represents 100 µm. **D.** Mean Viability of MDA-MB-231 in the device with 4 channels and 6

Similar to our experiments with the polystyrene beads, the high-speed camera videos of cell flows showed a constraint on the number of cancer cells that could occupy a given orbital within the vortex. Cancer cells entering an already occupied vortex would ‘knock out’ the existing one or get extruded out themselves. However, this number of trapped cancer cells can be scaled up by connecting multiple devices in series and parallel.

Finally, we also wanted to check if the cells subjected to the abovementioned forces for isolation were viable enough to be cultured for further assays. Experiments to check for the cellular viability using Trypan Blue (which selective stains dead cells) were carried out for MDA-MB-231 cells that were flowed in a device with 4 channels with 6 reservoirs. About 88.67% cells (with a standard deviation of 10.82%) were found to be alive after egressing from the device, implicating the potential compatibility of such isolation architectures for expansion and enrichment of CTCs (Fig. 5(D)).

## Conclusion

An architecture with inlet and outlet placed close to each other in orthogonal arrangement forms larger vortices at lower flow velocities. Owing to the combined interaction of forces arising from the turn-effect, i.e., shear-gradient lift-, centrifugal- and Dean-forces acting against an advective fluid drag force, the orthogonal vortex chip is capable of effective vortex trapping even at low flow velocities. This would not only help maintain the viability of cells, but also save the need of any special capacity pumps and apparatus that would be required to meet the high pressure flow rates involved in other such devices ^20, 29, 34, 35, 39, 40^. The major utilization would be in separating CTCs in a passive and marker-agnostic manner; which would be carried out at high throughput and yet would involve flow velocities that are reasonably low enough to not impair the viability of cells. Furthermore, the size-specific attractor-ring in the vortex and the ‘knocking-out’ phenomenon, along with the aspects of viability, purity and efficiency for the devices pertaining to CTC isolation are to be investigated in detail in future efforts.

## Materials and methods

### Sample Preparation

Polystyrene beads of size 15 µm, 20 µm (10.1% w/w) were purchased from Bangs Laboratories Inc. These were used to prepare particle sample solutions by diluting the particle suspensions in PBS or DI water, along with 0.1% v/v Tween 80 (Sigma-Aldrich) solution added to prevent particle aggregation. The mixture was mixed vigorously using a vortexer.

High concentration beads solution was used for experiments with CCD camera, consisting of bead concentration 1% v/v in DI water. This enabled ease in observing general particle trajectories and flow pathlines. Low concentration solution of bead and RBC mix was used for experiments with High-speed camera, containing bead concentration 0.09% v/v in 1X PBS, along with 1.66% v/v whole blood. This allowed observation of individual particle trajectories.

Cancer cell experiments were carried out with High-speed camera, with sample solution consisting of cancer cells (MDA-MB-231, MCF 7 and BT549) concentration ∼8×10^4^ cells/ml in 1X PBS along with 1.66% v/v whole blood.

### Cell Culture

Three human breast cancer cell lines MDA-MB-231, MCF-7 and BT-549 were cultured in Dulbecco’s Modified Eagle’s Medium supplemented with 10% FBS, respectively. The cells were cultured at 37 °C, 95% relative humidity and 5% CO_2_, which were then trypsinized (Trypsin-EDTA 1X, Sigma) and mixed with 1:1 media for use.

### Microfluidic Chip Fabrication

The microfluidic chip was fabricated using soft lithography techniques. Due to the high aspect ratio of the microfluidic channels, the mold for PDMS casting was made in silicon. A cleaned 4-inch silicon wafer was spin coated with positive photoresist AZ4562 (MicroChemicals GmbH) at 4000 rpm for 40 s to get a thickness of ∼6 µm. After a soft bake at 110°C for 1 min, the wafer was UV-exposed using Heidelberg µPG 501 to directly write the pattern. This was followed by development of the exposed photoresist using MF26A. The prepared wafer was then hard baked at 110°C for 4-5 mins followed by etching in deep reactive ion etching in DRIE (SPTS LPX Pegasus). Remaining photoresist was removed using oxygen plasma ashing processes in the DRIE tool. The height of the pattern was verified using Dektak XT surface profiler. The prepared wafer was then spin coated with Teflon^®^ (AF-2400) solution (DuPont) at 4000 □ rpm for 40 s to get a thickness of ∼170 nm. Spin coated Teflon was hard bake at 180 □ °C to evaporate the solvents. Teflon coating reduces adhesion of the casted PDMS and hence allows reuse of the master-mold multiple times.

PDMS (Sylgard 184, Dow Corning) mixed in a cross-linker to polymer ratio of 1:10 was cast on to the prepared mold and degassed in a vacuum desiccator. After curing at 60 °C for 4-5 hours, the PDMS devices were peeled off the wafer. Inlet and outlet holes were made using biopsy punch. The punched PDMS device along with a cleaned glass slide was treated with oxygen plasma (Harrick Plasma). The two activated surfaces were placed against each other to get PDMS-glass bonded microfluidic chips, which were then baked at 110°C for 1 h to get permanently bonded devices that do not leak at high pressures.

### Experimental Setup

Microparticle/cell samples were injected into the device from a plastic syringe using a New Era syringe pump (NE-8000). Owing to the high pressure and high flow rates involved, connectors were custom made using catheters, infant-feeding tubes, micropipette tips, and epoxy adhesive. (See Fig. S7) Imaging was carried out using either a CCD camera (IDS Imaging UI-3240CP-C) with a 3X lens, or a high-speed camera (FASTCAM Mini UX100, Photron) at 6400 fps coupled to microscope with 10X magnification lens. MATLAB R2019a was used to generate scatter plots through image processing techniques.

### Live Dead Analysis

A solution of MDA-MB-231 (3×10^6^ cells/ml) in PBS was used to perform the live dead experiment. The suspension was injected through the 3E design of 4 channels with 6 reservoirs each at an average inlet velocity of 1.3 m/s and collected. 0.4% Trypan blue (Himedia Laboratories Pvt. Ltd) was used in a dilution of 1:1 to count the live and dead cells in a Neubauer’s hemocytometer.

## Supporting information

ESI

## Authors’ Contributions

P. Seth proposed the presented idea. N. Rastogi, P. Seth, P. Sen and R. Bhat designed research. N. Rastogi performed microfluidic chip fabrication. N. Rastogi and P. Seth executed software simulation, preliminary tests and formal analysis. N. Rastogi validated and investigated experiments. N. Rastogi, P. Seth and P. Sen created experimental setup. N. Rastogi, P. Seth, P. Sen and R. Bhat wrote the manuscript. P. Sen and R. Bhat were responsible for supervision and funding acquisition.

## Acknowledgements

We thank Dharma Pally and Shahid Hussain for help with cell culture and experiments. The devices were fabricated and characterized in the National Nanofabrication Centre and Micro Nano Characterization Facility respectively. This work was supported by DBT Nanotechnology Funds. R.B would like to acknowledge funding support from the Department of Biotechnology, [BT/PR26526/GET/119/92/2017], SERB [ECR/2015/000280] and Institute of Eminence grant (IE/CARE-19-0319)].

