## Supplementary material for "Vortex chip incorporating an orthogonal turn for size-based isolation of circulating cells": ESI

*Part I – Supplementary Figures (S1-S7)*

*Part I – Supplementary Movies (M1-M5)*

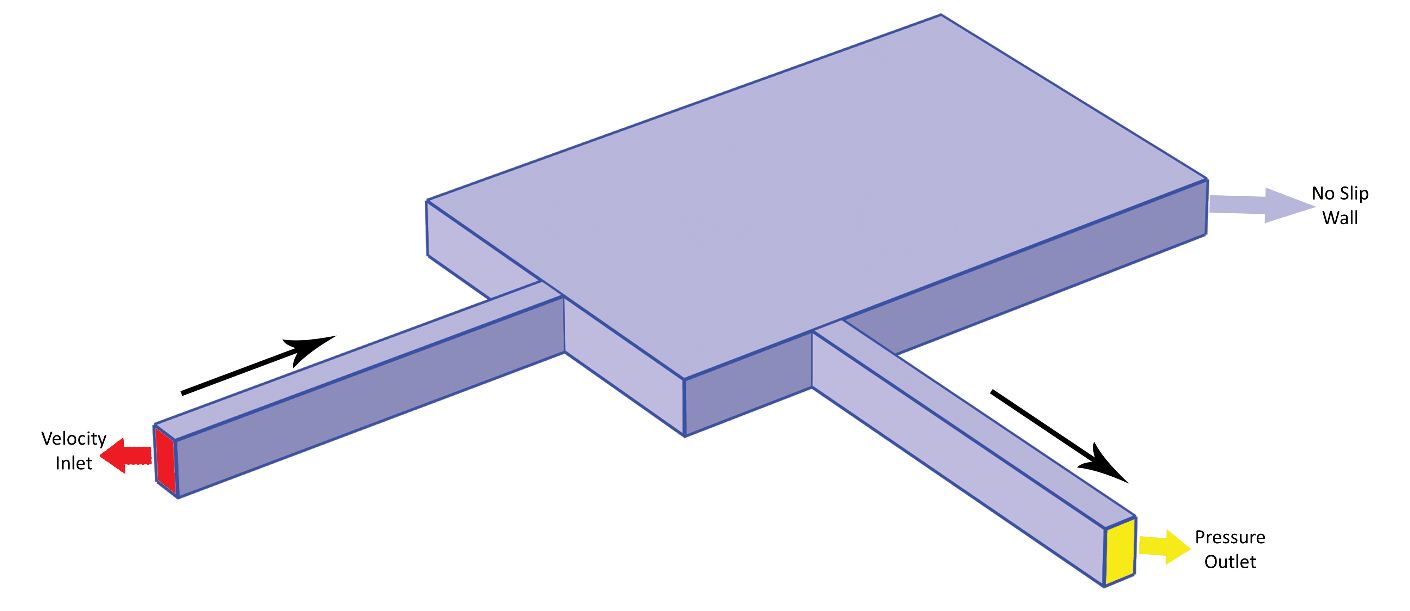

Figure S1: COMSOL Multiphysics boundary conditions.

Figure S2: **A** Certain designs exhibited highly irregular and unsteady trapping, where the particles either get trapped temporarily, or particles enter the vortex area and escape without getting trapped. Such designs were deemed unoptimal. The arched trajectory shows that the particles don’t follow the shortest path. A schematic representation of r_turn_ is also shown. **B** A map showing selected, unstable trapping and no trapping designs.

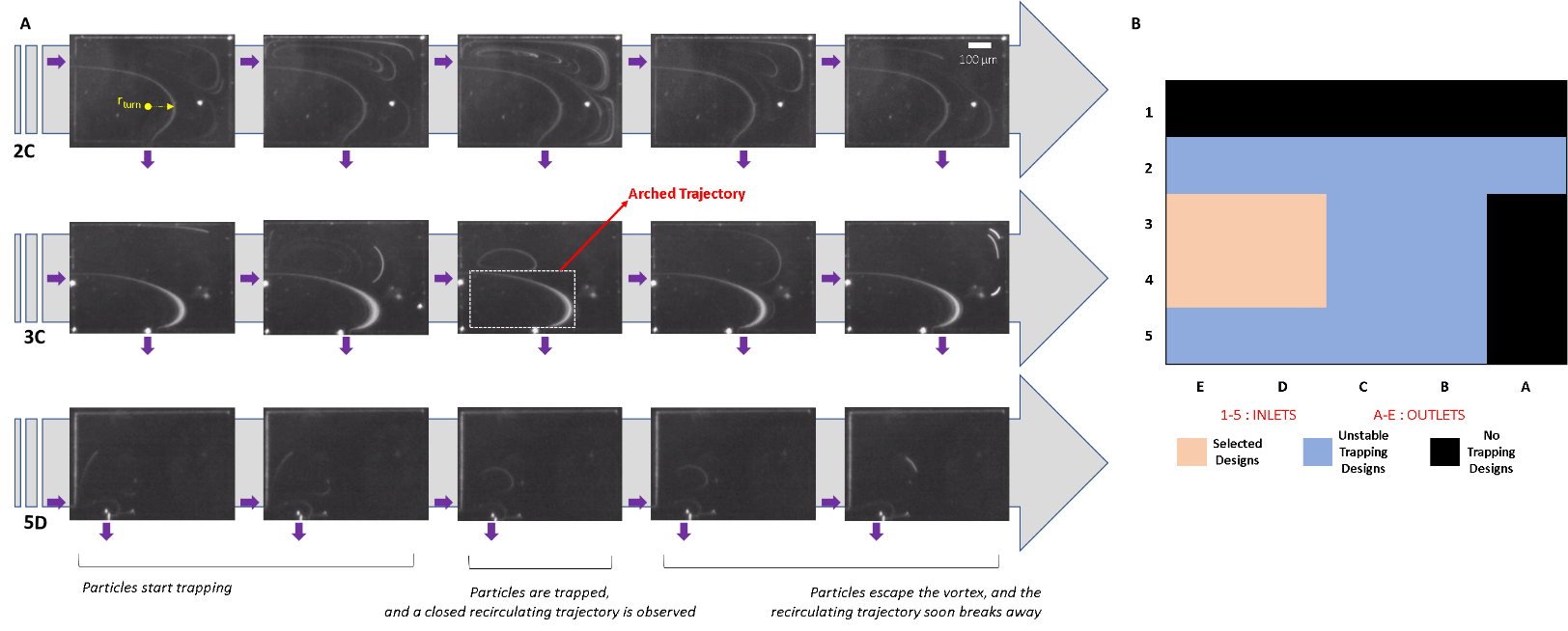

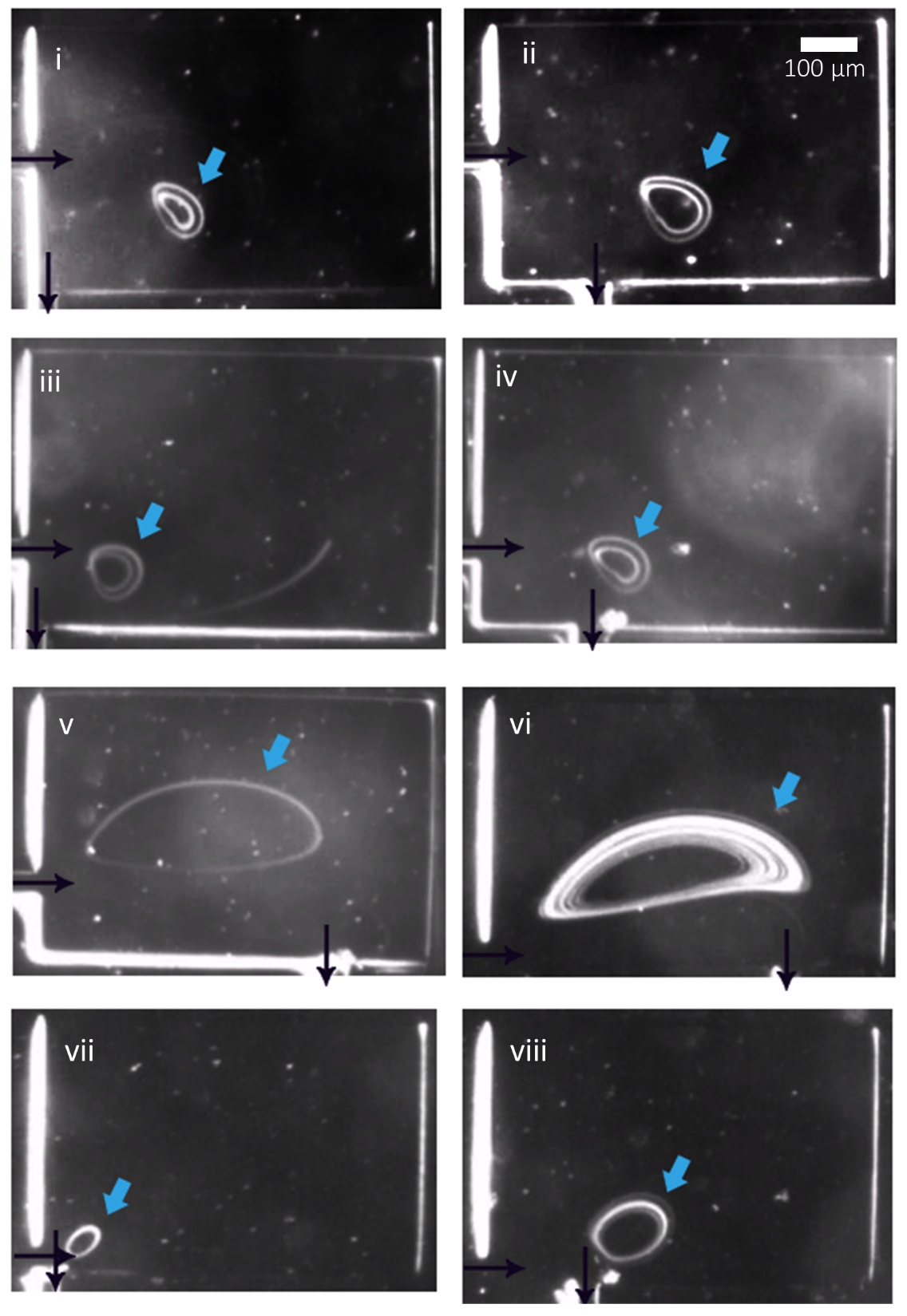

Figure S3: Vortex entrapment of 20µm beads in various designs. **i.** 3E **ii.** 3D **iii.** 4E **iv.** 4D **v.** 4B **vi.** 5B **vii.** 5E **viii.** 5D. There is an interplay between the flow-rate at which vortex-trapping is initiated, and the number of beads that are trapped (viz. the area of the trapping orbital or the ‘attractor-ring’). For instance, 5E traps beads at the lowest flow-velocity, but it’s attractor-ring is small in size. Scale bar represents 100µm. Arrows indicate inlet and outlet flow direction. Flows are at U = 0.4-2m/s.

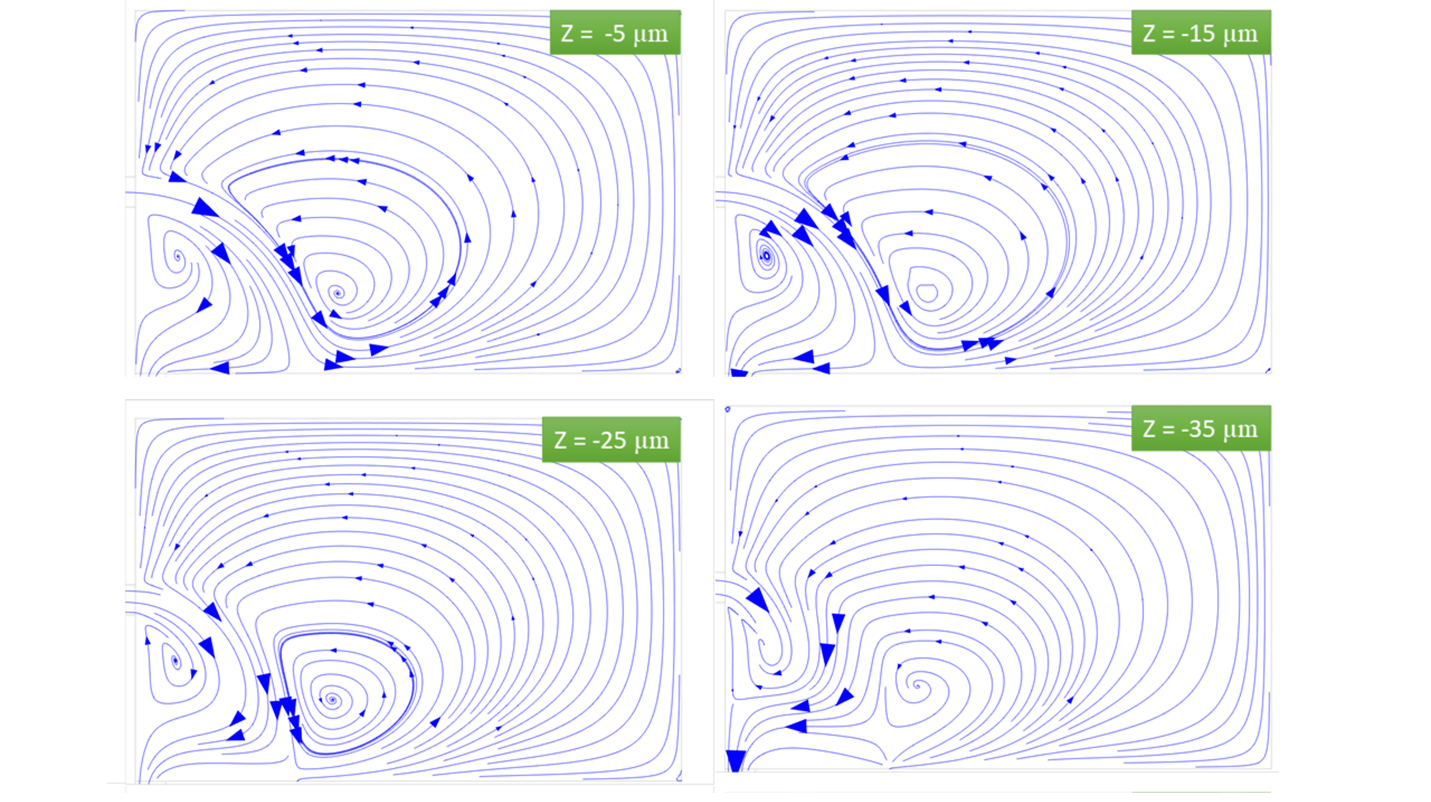

Figure S4: Distinct vortex profiles in planes at different depths (-35, -25, -15 and -5 µm) in the reservoir. (Central-plane is at Z=0.)

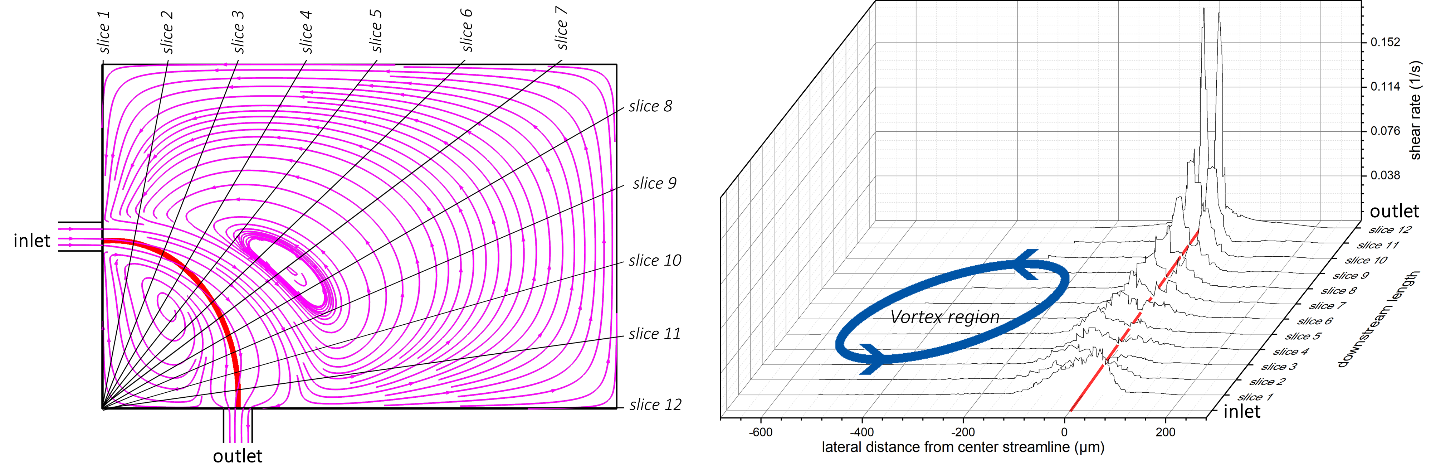

Figure S5: Shear rate distribution in 3D device at U=1.2m/s. Using COMSOL simulations, multiple slices were taken along the curvilinear trajectory to analyze the variation of shear rate values. The shear rate distribution is minimum towards the center of the vortex, indicating that the shear-gradient lift force which acts in the direction of decaying shear gradient would always push the particle towards the center of the vortex.

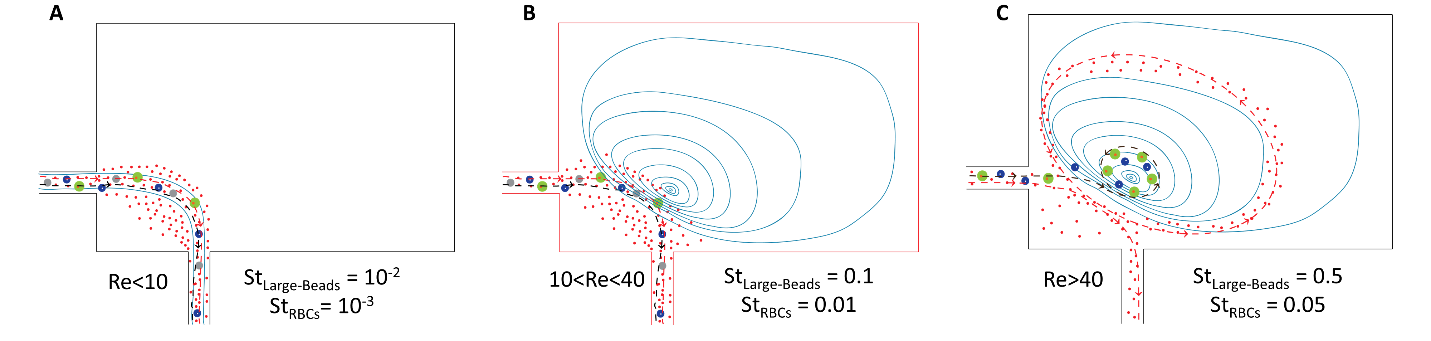

Figure S6: **(A-C)** Various regimes of the flow. **A.** At Re<10, flow is entirely laminar and creeping. St_larger_<<0.1 and St_rbc_<<0.1, consequently both larger particles and smaller RBCs enter and exit the reservoir along the flow stream. Particle trajectories are observed making a spindle. **B.** At 10<Re<40, vortex formation takes place. St_larger_<0.1 and St_rbc_<0.1, hence both larger particles and smaller RBCs still continue to move along the flow stream and no entrapment is observed. **C.** At Re>40, St_larger_>0.1 while St_rbc_<0.1. Hence, larger particles bearing larger inertia shoot through the flow-steam at the sudden orthogonal turn and transiting straight ahead get trapped in the steady trapping orbital close to the core of the vortex. Smaller particles, on the other hand, continue flowing along the stream or circulate in the unsteady outer region of the vortex and pass-by without trapping. An enclosed recirculating trajectory is observed for the larger particles.

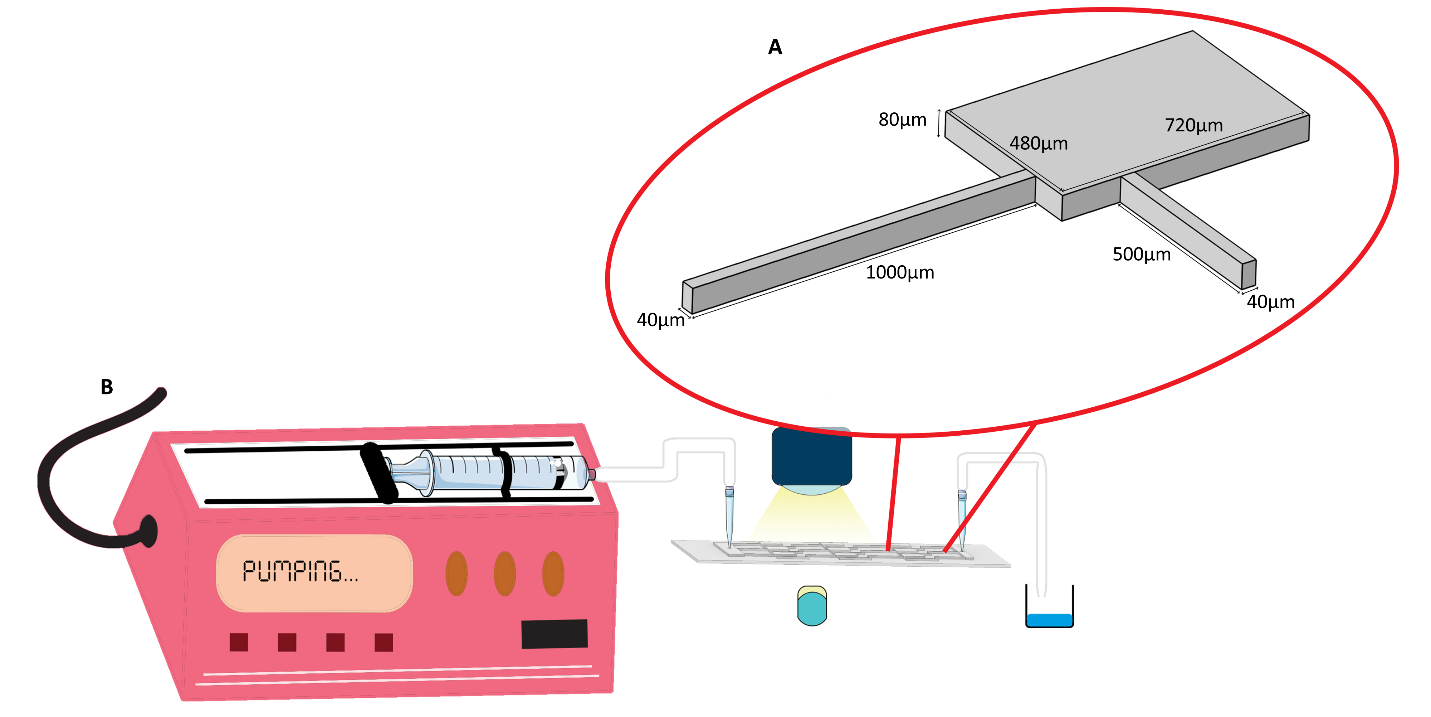

Figure S7: **A.** The dimensions of a non-linear vortex device. Shown here is a ‘4D’ device. The inlet and outlet channels are 1000µm×40µm×80µm and 500µm×40µm×80µm respectively, while the reservoir is 720µm×480µm×80µm. **B.** The experimental setup. Microparticle/cell samples are injected into the device from a plastic syringe using a syringe pump. Imaging is carried out using CCD or High-speed cameras coupled to microscope with 3X or 10X magnification lenses.

*Movie M1*: **A** In few high speed camera studies i. beads are observed to be trapped in different focal planes, indicating that the captured beads are recirculating in planes at different depths **ii.** A bead enters the reservoir and exits after traversing through the outer orbital without getting trapped. This confirms that the outer orbitals of the vortex do not enable stable trapping. **iii.** A new bead gets trapped with the already trapped beads when it enters the reservoir **iv.** A trapped bead gets knocked out of the attractor zone.

*Movie M2*: Size based vortex separation in a heterogenous mixture solution of RBCs and 20 µm beads. RBCs pass by undisturbed or follow about the unsteady orbits in the outermost zone of the vortex, while 20 µm beads get trapped in the steady trapping orbital close to the core of the vortex.

*Movie M3-M5*: Size-selective isolation of cancer cells from a heterogenous mixture with RBCs in PBS, for: (M3) MDA MB-231, (M4) MCF7, (M5) BT549.
